# Stratifying macrophages based on their infectious burden identifies novel host targets for intervention during Crohn’s disease associated adherent-invasive *Escherichia coli* infection

**DOI:** 10.1101/2024.01.10.575028

**Authors:** Xiang Li, John Cole, Diane Vaughan, Yinbo Xiao, Daniel Walker, Daniel M. Wall

## Abstract

Bacterial infection is a dynamic process resulting in a heterogenous population of infected and uninfected cells. These cells respond differently based on their bacterial load and duration of infection. In the case of infection of macrophages with Crohn’s disease (CD) associated adherent-invasive *Escherichia coli* (AIEC), understanding the drivers of pathogen success may allow targeting of cells where AIEC replicate to high levels. Here we show that stratifying immune cells based on their bacterial load identifies novel pathways and therapeutic targets not previously associated with AIEC when using a traditional homogeneous infected population approach. Using flow cytometry-based cell sorting we stratified cells into those with low or high intracellular pathogen loads, or those which were bystanders to infection. Immune cells transcriptomics revealed a diverse response to the varying levels of infection while pathway analysis identified novel intervention targets that were directly related to increasing intracellular AIEC numbers. Chemical inhibition of identified targets reduced AIEC intracellular replication or inhibited secretion of tumour necrosis factor alpha (TNFα), a key cytokine associated with AIEC infection. Our results have identified new avenues of intervention in AIEC infection that may also be applicable to CD through the repurposing of already available inhibitors. Additionally, they highlight the applicability of immune cell stratification post-infection as an effective approach for the study of microbial pathogens.

## Introduction

Infection is a dynamic process with a highly heterogenous population of host cells infected to varying degrees by infiltrating microorganisms. These differing microbial loads can lead to a variety of outcomes for the infected cells and a heterogeneity in host responses. Replicating the dynamics of infection using either *in vivo* or *in vitro* models of disease is challenging, but these models have proved highly useful tools in understanding specific aspects of infection. While heterogeneity fundamentally underlies *in vivo* models of disease, *in vitro* models by design are often based on the interaction between a single pathogen-and a particular host cell type in a more controlled environment. This reduction in complexity has clarified aspects of the host or microbial response to infection, confirming or raising hypotheses for later testing in more complex models.

*In vitro* models of bacterial infection often require high multiplicities of infection (MOIs) to ensure a bacterial intracellular burden high enough to enable host-pathogen dynamics to proceed in a measurable way over time. While MOIs into the hundreds are common, these rarely result in homogenous infection by, or phagocytosis of, all bacteria present within the system. What results is a mixture of sub-populations with varying degrees of infectious load, with either no bacterial infection having occurred, low levels of intracellular bacteria or a high intracellular bacterial load. Yet these diverse sub-populations have traditionally been studied as a single homogenous population, leading to the potential loss of information critical to understanding the infection process. For example, there may be contrasting outcomes in immune cells where bacteria are overcome in some cells, while actively replicating intracellularly in others, yet the basis of these outcomes are generally not investigated in *in vitro* models.

Adherent-invasive *Escherichia coli* (AIEC) is a pathobiont isolated in increased frequency from the intestine of CD patients relative to healthy controls (Darfeuille-Michaud *et al*., 1998; Martinez-Medina *et al*., 2009; Nadalian *et al*., 2021). CD is a multifactorial disease with genetic susceptibility, dietary factors and microorganisms all playing a role in disease pathogenesis. Rising incidence, the increasingly young age of onset, and incurability of the disease mean that as well as reducing quality of life, CD is a significant burden on health care systems across the world (Bassi *et al*., 2004; Rao *et al*., 2017). While genetic susceptibilities linked to CD are well defined, specific defects in autophagy and protection against intracellular bacteria have not explained why bacteria such as AIEC are found with increasing prevalence. AIEC lacks many of the classic virulence factors associated with *E. coli* pathotypes and its persistence in the CD gut is likely mediated via metabolic success and adaption to the conditions in the inflamed CD gut (Ormsby *et al*., 2019, 2020; Cho *et al*., 2022; Sugihara *et al*., 2022). A hall mark of AIEC infection is replication to high levels within infected macrophages, where it can stall cell death pathways, a likely contributory factor in granuloma formation (Meconi *et al*., 2007; Dunne *et al*., 2013). With a paucity of information regarding the key drivers for the success of infection in the host-pathogen relationship, the treatment of AIEC infection in the context of CD has proved challenging, although recent progress has been made (Boucher and Barnich, 2022; Douadi *et al*., 2022; Gerner *et al*., 2022; Titécat *et al*., 2022). However, while AIEC replicates and persists to high levels in some infected macrophages this does not occur in all infected cells. Here we show that the population of AIEC infected macrophages is highly heterogenous, and this is reflected in the vastly different responses of cells to infection. While many cells remain uninfected, or have overcome AIEC infection, these cells remain within the studied *in vitro* population contributing to outputs and thus disguising the response to infection in cells where AIEC are actively infecting. By stratifying macrophages based on their infectious load, we identified host pathways significantly differentially expressed in direct response to infectious burden, information lost when treating cells as a single homogenous population. By inhibiting the identified differentially expressed pathways, which had not previously been linked to AIEC infection, we could block bacterial intracellular replication and release of the cytokine tumour necrosis factor alpha (TNFα), known to be a critical driver of inflammation in both AIEC infection and CD.

Our approach here shows that stratifying immune cells based on their bacterial load identifies novel pathways and therapeutic targets not detected using a traditional homogenous population approach. By focusing on host responses directly linked to bacterial success in cells where they are overwhelming the immune response, a more relevant and useful understanding of the complexities of infection can be gained.

## Materials and Methods

### Cell culture and infection

RAW 264.7 cells were seeded at a density of 2 x 10^5^ cells/ml into a T75 flask with 15 ml of Roswell Park Memorial Institute (RPMI) media (supplemented with 3% foetal bovine serum (FBS), penicillin/streptomycin and L-glutamate). Six hours post-cell seeding, RAW 264.7 cells were treated with 100 ng/ml of lipopolysaccharide (LPS) and incubated overnight. RAW 264.7 cells in RPMI-1640 with 3% FBS without antibiotics, were then infected with LF82 carrying the p*rpsM*GFP plasmid (LF82*rpsM*GFP) at a multiplicity of infection (MOI) of 100 for 1 hour (Li *et al*., 2022). Post-infection, extracellular bacteria were removed by washing with fresh RPMI media (3% FBS) containing 50 μg/ml of gentamicin and the media was replaced with fresh RPMI media (3% FBS, 50 μg/ml gentamicin). After 24 hours, cells were harvested using cell scrapers. Suspended cells were washed and maintained in fluorescence associated cell sorting (FACS) solution (2% FBS in phosphate buffered saline (PBS)). The viability of cell cultures was assessed using 7-aminoactinomycin D (7-AAD) viability staining solution at a final concentration of 0.25 μg/million cells. For each experiment four independent biological replicates were carried out with 4 technical replicates within each.

### Sorting of infected cells

Flow cytometry was performed on a BD FACSAria IIU with BD FACSDiva software version 9.0.1 (BD Biosciences, Franklin Lakes, NJ) paired with FlowJo Version 6.3.2 analysis software (Tree Star Inc., Ashland, OR). The instrument has not been altered and has a fixed- alignment cuvette flow cell and four-laser base configuration. Each sample was first examined using forward scatter (FSC) versus side scatter (SSC). Green fluorescent protein (GFP) was excited by a 488 nm, 20 mW Coherent laser and the emissions detected with a 530/30 bandpass filter set while 7-AAD was excited by a 561 nm, 50 mW Coherent laser, and the emissions picked up in the 660/20 Bandpass filter set. Based on measurements obtained from the analysis of 10,000 events for each samples, gating strategies were established for the selection of cells of interest using FSC, SSC, and fluorescence emission properties (Figure S1). Actual cells were easily distinguished from debris by gating on FSC and SSC. 7AAD was used to identify live/dead cells. In LF82 p*rpsM*GFP infected RAW 264.7 cells, living cells were gated based on their lack of 7AAD staining. A gating strategy was then established for the three populations of infected cells by determining their GFP fluorescence intensity. The identification of different intracellular bacterial burdens as *No*, *Low* and *High*, were used to sort the cells into three separate populations, representing cells with no bacteria (*No*), cells with less than 5 bacteria (*Low*) and cells with more than 5 bacteria (*High*). The control group cells, where bacteria had not been added, were sorted in the same number as for the other three groups. Data was acquired for each population for 80,000 cells. To simplify the description of the four groups of cells in the following text, the terms “*Control*”, “*No*”, “*Low*” and “*High*” will be used to indicate their infection status. Sorted cells were collected into 1.5 ml microfuge tubes containing 800 μl of RNAlater solution (Invitrogen AM7020) stopping cellular transcriptional changes. RNA from four independent biological repeats was collected and kept at -80°C until RNA was extracted.

### RNA isolation

RNA was extracted using an RNeasy PowerMicrobiome Kit (QIAGEN, 26000-50) using the manufacturer’s protocol. RNA extracts were kept at -80°C. Both quantity and quality of RNA were assessed by using an Agilent 2100 Bioanalyzer (Agilent Technologies). RNA yields ranged from 3.47 to 18.6 ng/μl. RNA integrity numbers (RIN) of a sample are generated by the 2100 Bioanalyzer to indicate the level of degradation and have been shown to predict gene expression suitability reliably. RIN scores ranged from 8.7 to 10, indicating high-quality RNA suitable for gene expression analysis by RNA sequencing (RNA-seq) (Fleige and Pfaffl, 2006).

### Library construction, RNA-seq, and bioinformatics

At least 10 ng of RNA was isolated per sample and provided to Glasgow Polyomics (University of Glasgow) for RNA-seq, the generation of cDNA, sequencing, and bioinformatics. The cDNA libraries were created using the Quantseq (FWD) kit from Lexogen. The kit creates a library from the polyA end of transcripts, creating fragments terminating in the polyA sequence and sequencing towards this. The libraries were sequenced at 75bp, paired end, to a mean depth of 10 million reads per sample, using an illumine (NextSeq 2000). The data was quality controlled and aligned using Galaxy (server: http://antioch.tcrc.gla.ac.uk/). Firstly, read quality was explored using FastQC, then trimmed using Trimmomatic (Bolger, Lohse and Usadel, 2014), under default settings. Reads were mapped to the reference genome (GRCm39) and transcriptome (v110) using Hisat2 (Kim *et al*., 2019), under default settings. Read counts were produced using HTseq-count (Anders, Pyl and Huber, 2015), which were then normalised, and pairwise differential expression calculated using DESeq2 (Team RC., 2014). Searchlight (Cole *et al*., 2021) was used to explore and visualise the data. Each pertinent pairwise comparison was entered as a DE workflow, with (adjusted p < 0.05 and absolute log2fold > 1). A single MDE workflow was used combining each of No, Low and High vs Control comparisons. For the pathway analysis the KEGG and GO pathway databases were used with (adjusted p < 0.05).

## Results

RAW 264.7 cells that had been incubated with LF82*rpsm*GFP, or control uninfected cells, were subjected to flow cytometry-based cell sorting to isolate cells based on the intensity of green fluorescence and the number of intracellular bacteria enumerated by colony formation unit (CFU) counts. The experimental procedure is outlined in Figure 1a. Based on fluorescence intensity, RAW264.7 cells co-incubated with LF82*rpsm*GFP led to 3 distinct populations of cells (each population was at least 80,000 cells) (Figure 1b); those that remained uninfected despite being in proximity to LF82*rpsm*GFP (*No*), those with a bacterial load with an average of 1-2 bacteria per cell (*Low*) and those with approximately 7 bacteria per cell (*High*) (Figure 1c). The control uninfected cells had no contact with LF82*rpsm*GFP (*Control*).

**Figure 1:**
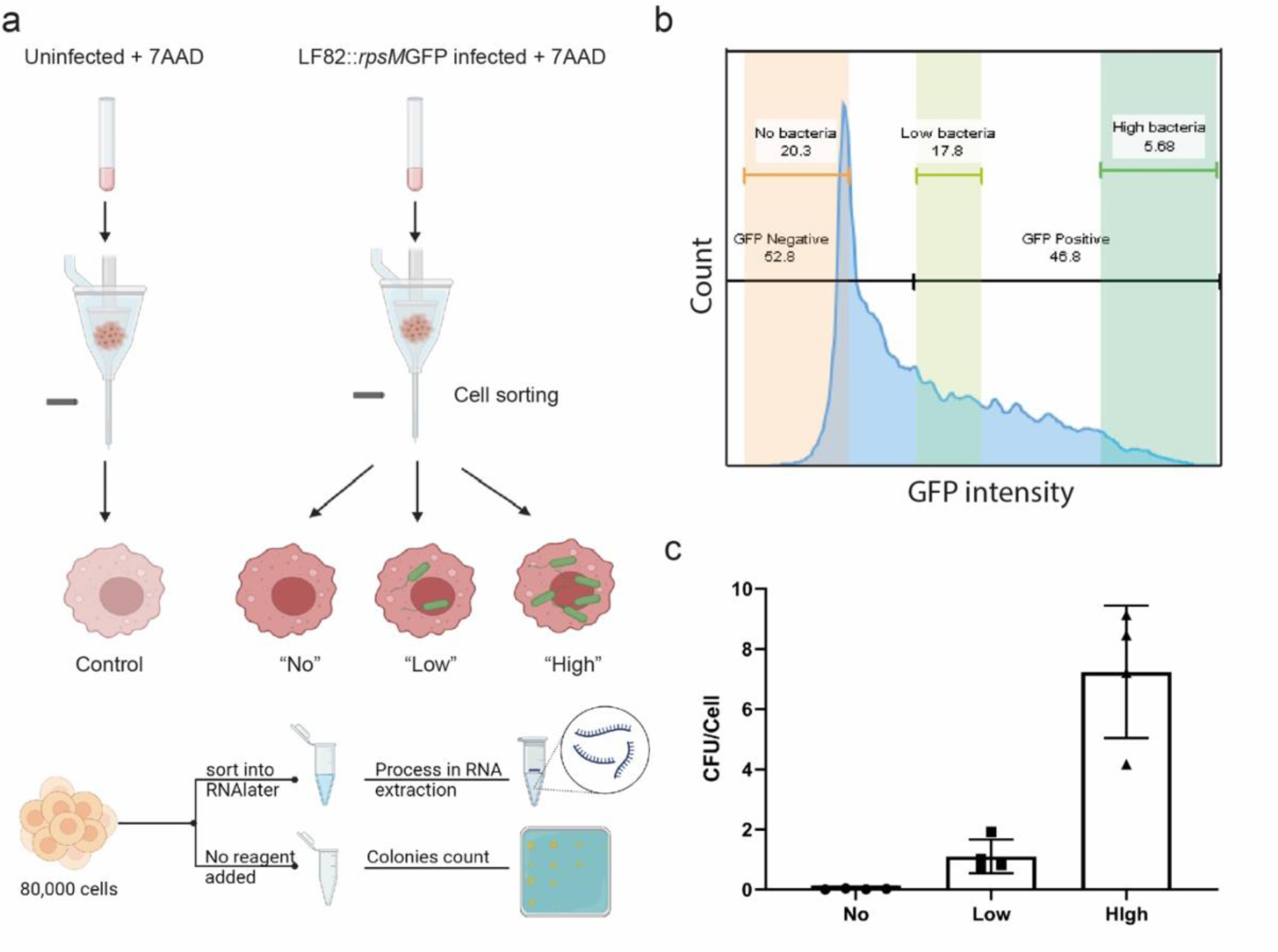
Macrophage sub-populations sorted by FACS and confirmation of intracellular bacteria number by traditional visible colony count. (a) Schematic overview of the process of sorting RAW 264.7 cells for RNA-seq and viable count analysis. There were 4 independent biological repeats, each repeat includes two sorts: one was sorted into an RNAlater solution, enabling later RNA extraction; another sort was used for confirming the number of intracellular bacteria. (b) Three populations of cells were determined according to GFP intensity. (c) The number of intracellular bacteria from different populations was calculated after their recovery by plating it onto LB agar plate and CFU counting.

RNA extraction was carried out from sorted cells and differential expression analysis was undertaken. Each population of *Control*, *No*, *Low* and *High* RAW 264.7 cells were compared to each other to identify differentially expressed genes (DEGs) between each group (Figure 2). Principle component analysis (PCA) clearly showed the cells from the infected population clustering together and away from uninfected cells as expected (Figure 2a). While it was clear from the resulting heatmap and a count of significant DEGs that the response in *control* cells was significantly changed in comparison to cells in proximity to, or with intracellular LF82*rpsm*GFP, it was also noted that there was a significant change in response between each of the *No*, *Low* and *High* sub-populations of cells (Figure 2b-c). Comparing DEGs between *Control* cells and those where LF82*rpsm*GFP were present (*No*, *Low* and *High*), 28.8% of significant DEGs were common to all cells from this population (Figure 3a).

**Figure 2:**
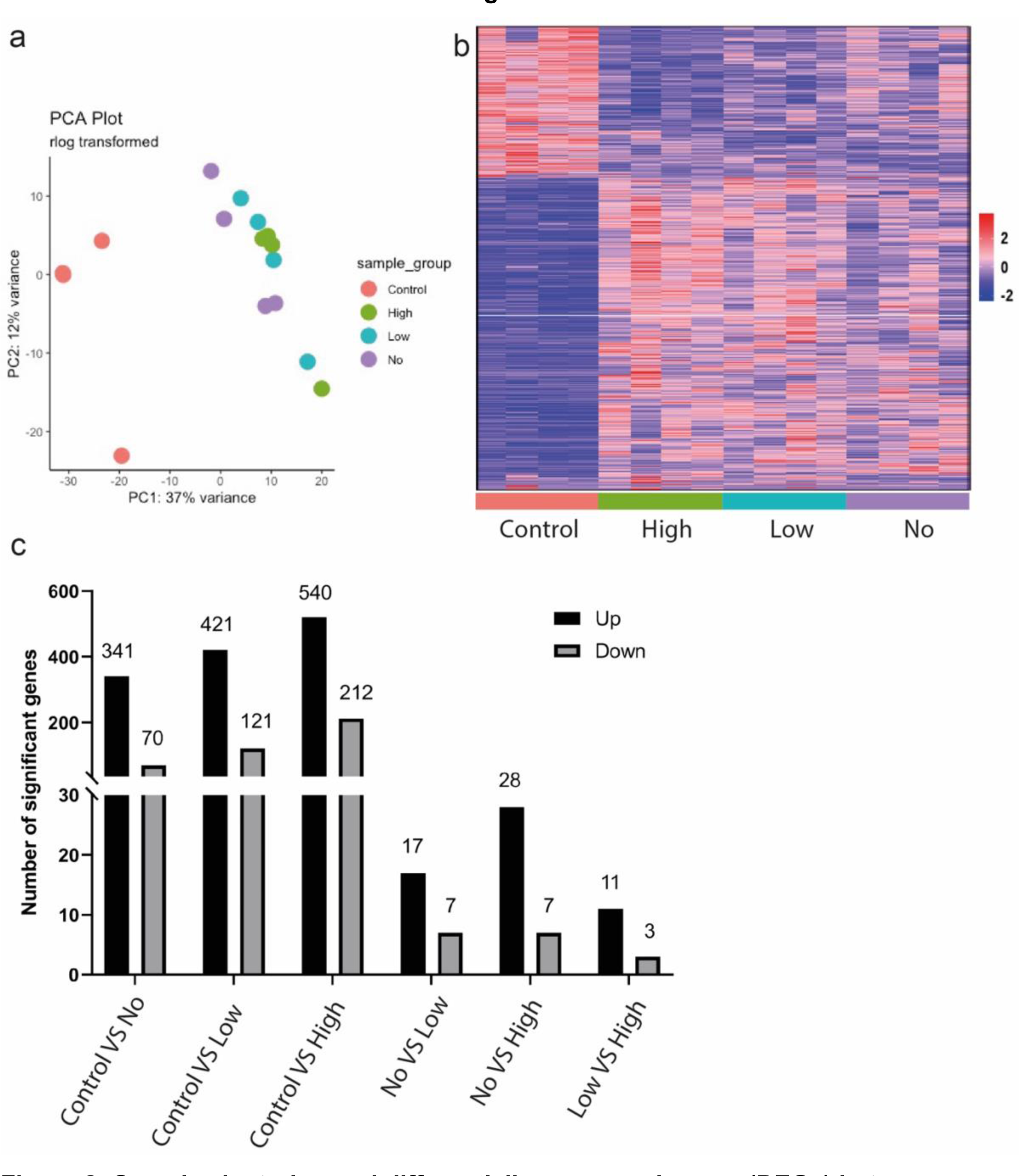
Sample clustering and differentially expressed genes (DEGs) between different macrophage populations with differing bacterial burdens. (**a**) Principal component analysis (PCA) of expression data, the first two components. Dots represent replicates and are coloured by condition (red=Control, green=High, blue=Low, purple=No). The % variance is given. (**b**) Expression heatmap of all DEGs (adjusted p < 0.05 and absolute log2fold > 1) in any of 6 comparisons (*Control* vs *No*, *Control* vs *Low*, *Control* vs *High*, *No* vs *Low*, *No* vs *High*, *Low* vs *High*). Axis are hierarchically clustered. Expression values are per gene Z-scores with low=blue and high=red.

**Figure 3:**
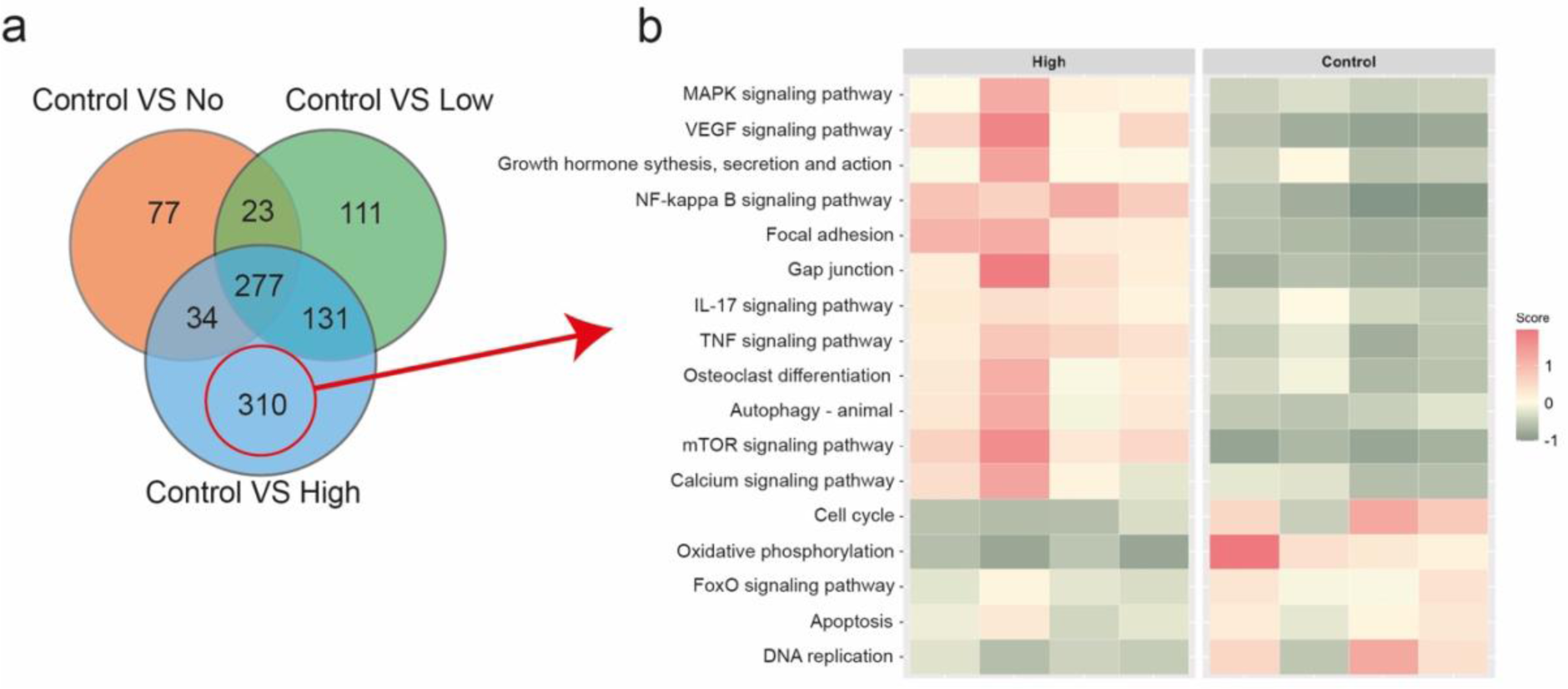
Characteristics of unique DEGs in the comparison of *Control* vs *High*. (**a**) Venn diagram showing the number of overlapped or unique DEGs (adjusted p < 0.05 and absolute log2fold > 1) in the three comparisons: *Control* vs *No*, *Control* vs *Low* and *Control* vs *High*. There are 310 unique DEGs in the comparison of Control vs High. (**b**) Heatmap of enriched KEGG pathways (adjusted p < 0.05) for the 310 unique genes in (a). The heatmap shows mean expression across all genes in the enriched pathways, with the rows being pathways and columns individual samples. Red indicates relative pathway activation and *green* represents relative pathway suppression.

However, there were also significant changes in responses between the cells in the infected population with 32.2% of DEGs between the infected and uninfected populations unique to the *High* group, 11.5% of DEGs unique to the *Low* group, and 8% of DEGs unique to *No* group (Fig. 3a). This pointed towards a clearly heterogenous population with cells that were uninfected but bystanders to infection of other cells (*No* group) having their own unique response, acting in a more similar fashion to infected rather than uninfected cells. Pathway analysis was conducted on the 32.2% of DEGs unique to the *High* and *Control* sub- populations. The outcome clearly identified several pathways associated with the immune response that were activated in the *High* group, including the nuclear factor NF-kappa B (NF- κB) pathway, while pathways related to the cell cycle were inhibited in the *High* group (Fig 3b).

Analysis of cytokine gene expression again clearly indicated differences between sub- populations within the total infected population. While expression of many cytokine-related genes was increased within the infected population, the bystander cells without bacteria (*No* group) were noted to have lower expression of several related genes (e.g. TNFα receptor: *Tnfrsf1b*), while other genes were expressed at similar levels to those cells with *High* bacterial load (e.g. *Tnf*; Figure 4). Therefore, these bystander cells, were clearly contributing to inflammation through cytokine production but were not being influenced to the same extent by circulating cytokines such as TNFα.

**Figure 4:**
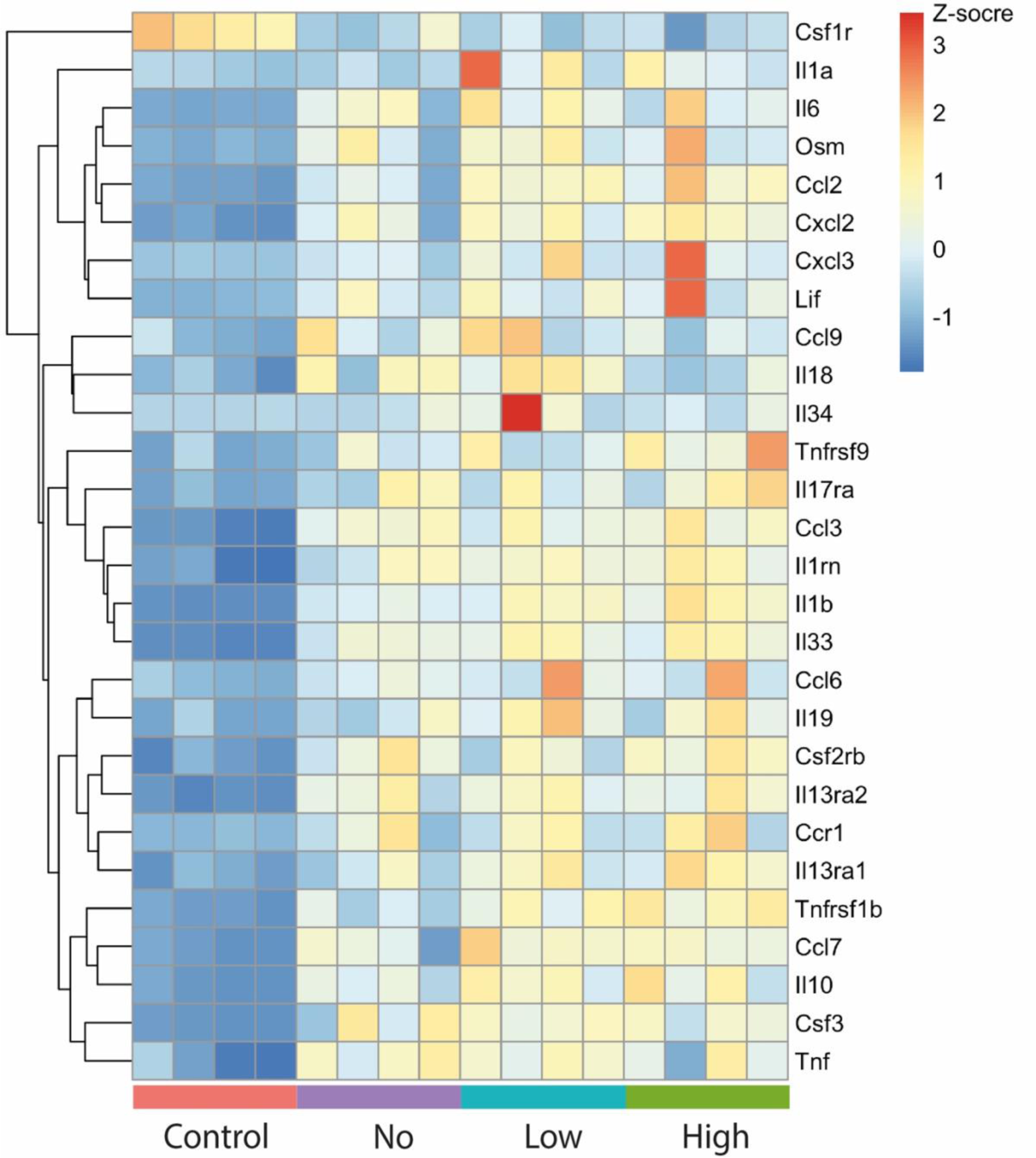
Heatmap of changes in gene expression levels of cytokine and chemokine genes in three groups infected with LF82 (No, Low and High) alongside the Control uninfected group. Heatmap of differentially expressed cytokines (adjusted p < 0.05 and absolute log2fold > 1) between each of No, Low and High vs Control. Rows represent cytokines and columns samples. The y-axis is hierarchically clustered. Expression values are per gene Z-scores with low=blue and high=red.

Having determined both intracellular LF82*rpsm*GFP numbers and gene expression in response to intracellular bacterial load, we used this data to determine signatures of host gene expression in response to LF82*rpsm*GFP and identify host pathways that were expressed or repressed in response to infectious burden (Figure 5). As pathway analysis was not possible in the context of three pairwise comparisons (*No* vs *Low*, *No* vs *High* and *Low* vs *High*) due to their low number of significant DEGs, this approach of determining *signatures of infection* allowed us to extract valuable information related to infection status and drivers of increased infectious burden. We could therefore move past the simple comparison of DEGs in the context of infected versus uninfected cells and examine significant DEGs in the context of the heterogenous AIEC infected population. Two signatures of infection were tested, *Signature 1* selected for significantly increased DEGs that increased stepwise in direct response to increasing LF82*rpsm*GFP burden. 516 genes fitting these criteria (Fig. 5a). *Signature 2* selected for significantly increased DEGs that had an inverse relationship with intracellular LF82*rpsm*GFP burden, their expression decreasing as bacterial burden increased, 222 genes fitted these criteria (Fig. 5b). *Signature 1* clearly showed that, as bacterial numbers increased, there was a corresponding increase in pathways related to inflammation, chemotaxis and response to bacterial stimuli (Fig. 5a, Table 1). *Signature 2* showed that increasing intracellular LF82*rpsm*GFP load was inversely related to pathways for RNA metabolism, ribosome assembly and cell differentiation, all of which were significantly lower in cells with higher intracellular bacterial loads (Fig. 5b, Table 2).

**Figure 5:**
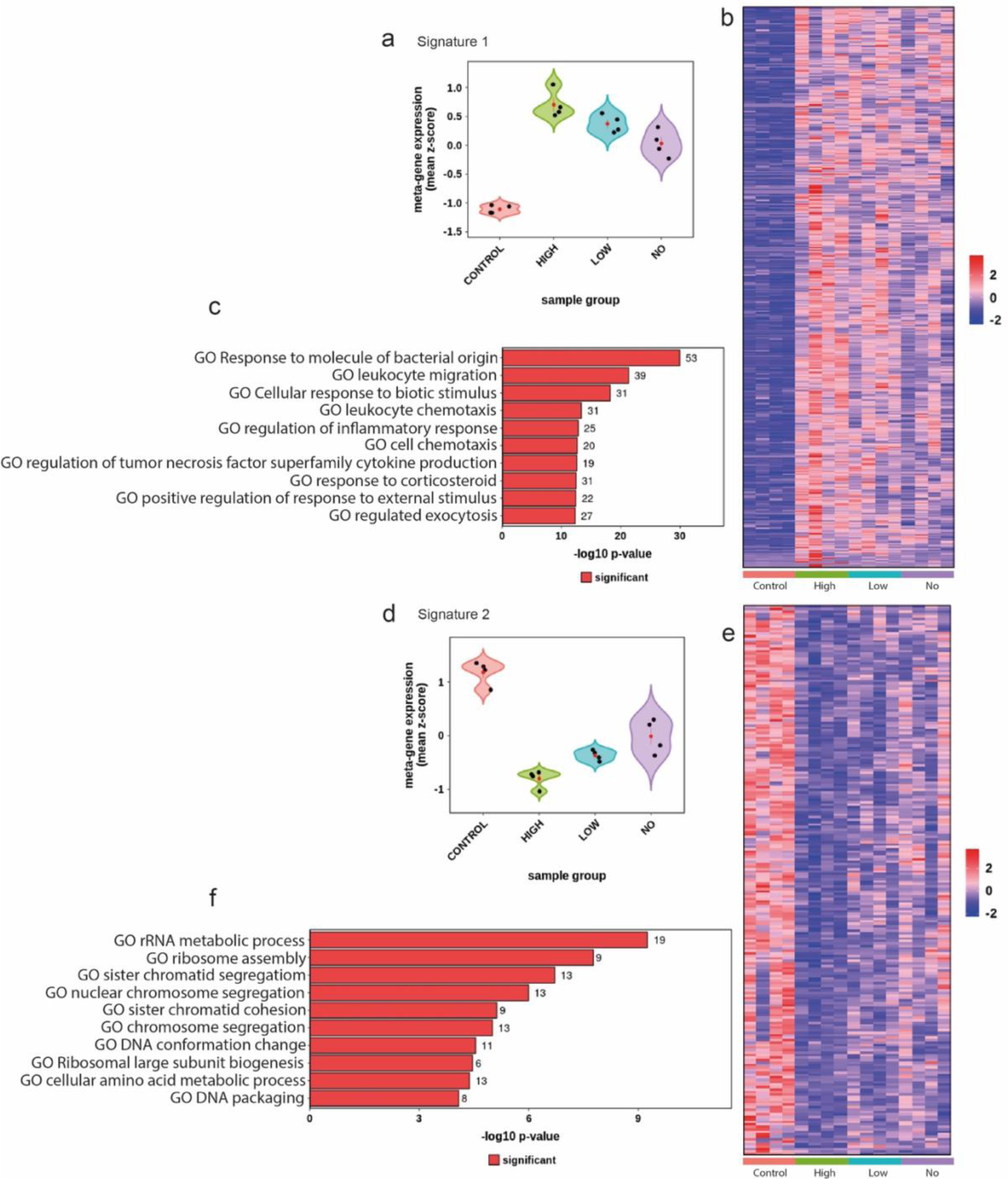
Signature gene expression among 4 populations and their relevant enriched GO-BP pathways. (**a**) Signature analysis for genes that are elevated (adjusted p < 0.05 and log2fold > 1) in all groups (*No, Low, High*) vs *control*. Showing: (left) metagene violin plot, with the mean expression z-score on the y-axis and group on the y-axis; (right) expression heatmap for all genes in the signature, showing genes by row and samples by column. The y-axis is hierarchically clustered. Expression values are per gene Z-scores with low=blue and high=red; (bottom) ten most enriched GO biological processes (p.adj < 0.05) for the signature genes. Showing the -log10p adjust value on the x axis and the number of DEGs in each enriched pathway as the data label. **(b)** as (a) however for the genes that are downregulated (adjusted p < 0.05 log2fold < -1) in all groups (*No, Low, High*) vs *control*.

**Table 1:**
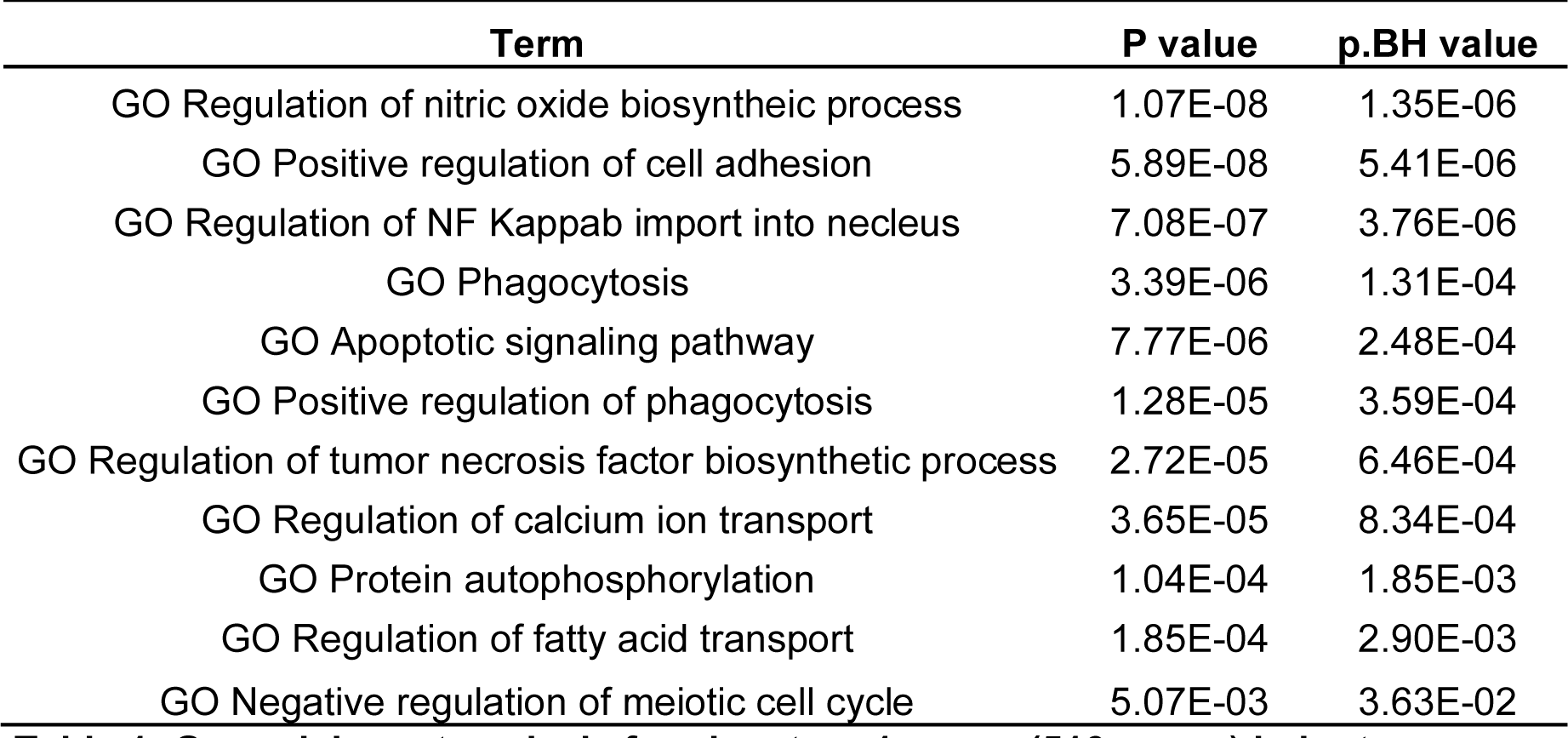
Go enrichment analysis for signature 1 genes (516 genes) in bp terms.

| Term | P value | p.BH value |
| --- | --- | --- |
| GO Regulation of nitric oxide biosynthetic process | 1.07E-08 | 1.35E-06 |
| GO Positive regulation of cell adhesion | 5.89E-08 | 5.41E-06 |
| GO Regulation of NF Kappab import into nucleus | 7.08E-07 | 3.76E-06 |
| GO Phagocytosis | 3.39E-06 | 1.31E-04 |
| GO Apoptotic signaling pathway | 7.77E-06 | 2.48E-04 |
| GO Positive regulation of phagocytosis | 1.28E-05 | 3.59E-04 |
| GO Regulation of tumor necrosis factor biosynthetic process | 2.72E-05 | 6.46E-04 |
| GO Regulation of calcium ion transport | 3.65E-05 | 8.34E-04 |
| GO Protein autophosphorylation | 1.04E-04 | 1.85E-03 |
| GO Regulation of fatty acid transport | 1.85E-04 | 2.90E-03 |
| GO Negative regulation of meiotic cell cycle | 5.07E-03 | 3.63E-02 |

**Table 2:** GO enrichment analysis for signature 2 genes (222 genes) in BP terms.

| Term | P value | p.BH value |
| --- | --- | --- |
| GO rRNA metabolic process | 8.96E-10 | 3.62E-06 |
| GO Ribosome assembly | 2.38E-08 | 4.81E-05 |
| GO DNA Conformation change | 2.18E-05 | 1.26E-02 |
| GO DNA packaging | 6.05E-05 | 2.44E-02 |
| GO Cellular amino acid metabolic process | 6.87E-05 | 2.52E-02 |
| GO Alpha amino acid metabolic process | 1.36E-04 | 4.30E-02 |

To further investigate the importance of these pathways to LF82 infection several significant DEGs were selected from the highlighted *Signature 1* pathways with each DEG showing the Signature 1 stepwise increase in expression correlating with increased intracellular LF82*rpsm*GFP burden (Fig. 6, Table 3). Chemical inhibitors were identified for a number of these *Signature 1* gene products that could be used to test their role in LF82*rpsm*GFP infection; ST034307 for Adcy1, clomipramine for Itch, trametinib for Map2k1, necrosulforamide for Mlkl, and GSK2636771 for Pik3cb (Table 4). Importantly none of the selected inhibitors had previously been tested in the context of either AIEC infection or CD. RAW 264.7 cells were again exposed to LF82*rpsm*GFP and phagocytosis was allowed to occur prior to treatment to prevent any inhibition of pathogen uptake influencing the results. Each of ST034307, clomipramine and GSK2636771 were seen to influence intracellular bacterial burden at 24 hours post infection (hpi) (Fig. 7ab and 7e). While the inhibitors were not toxic to bacteria during growth, it was clear that at high concentrations certain inhibitors were cytotoxic to cells (Fig. S2). So, while ST034307 inhibition of Adcy1 function significantly decreased intracellular LF82*rpsm*GFP at 24 hpi, it was seen to induce increases in cytotoxicity when used at the effective 10 μM concentration, with this increase in cytotoxicity becoming significant upon infection. Clomipramine was determined to exhibit the most significant effects, reducing intracellular bacterial burden 3 log-fold (Fig. 7b). While clomipramine exhibited some cytotoxicity this was at a higher concentration than those that reduced intracellular bacterial load (Fig. S2). However, to rule out any cytotoxic effects on bacterial load, a reduced 1 μM concentration of clomipramine was tested over a longer time course (72 hpi) and the intracellular bacterial burden of live cells determined. Clomipramine was observed to significantly reduce both the number of infected cells and the intracellular bacterial burden in the remaining infected cells (Fig. 8). This effect of clomipramine was observed at 24 hpi (Fig. 8a) and continued over 48 (Fig. 8b) and 72 hpi (Fig. 8c) with the number of infected cells reducing by half and the number of cells with a *High* bacterial burden reducing by over two-thirds. No changes in bacterial burden were observed with the other inhibitors.

**Figure 6:**
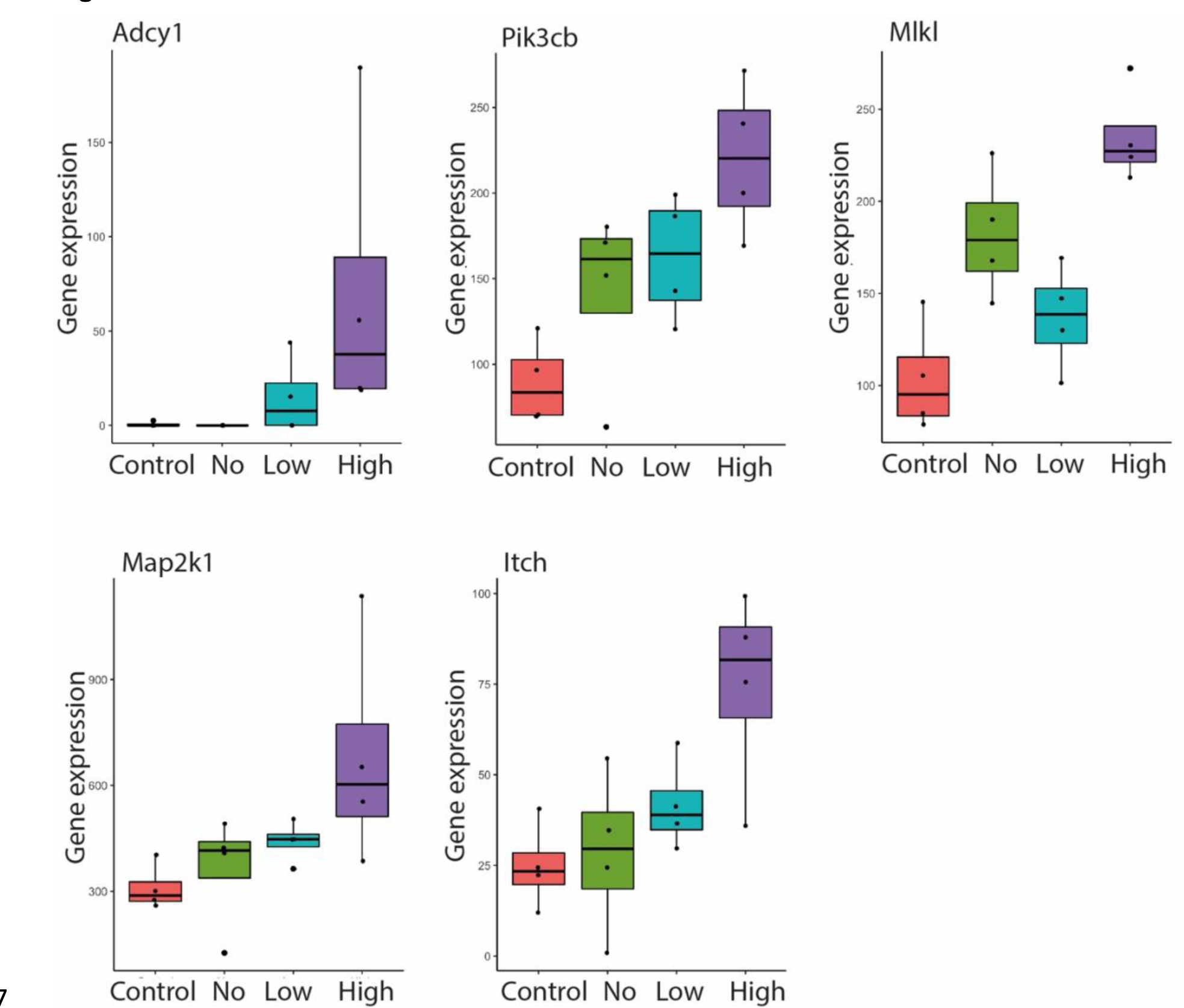
Gene expression levels of five candidate host DEGs selected for further testing. Genes *Adcy1*, *Pik3cb*, *Mlkl*, *Map2k1* and *Itch* were selected from the signature 1 gene list involved in pathways; cell-cell adhesion, TNF signalling, necrotic cell death, MAPK pathways and NF-κB. Boxplots show expression of genes of interest in four groups: *Control* in red, *No* in green, *Low* in blue and *High* in purple. Black dots denote individual samples. Error bars represent SEM.

**Figure 7:**
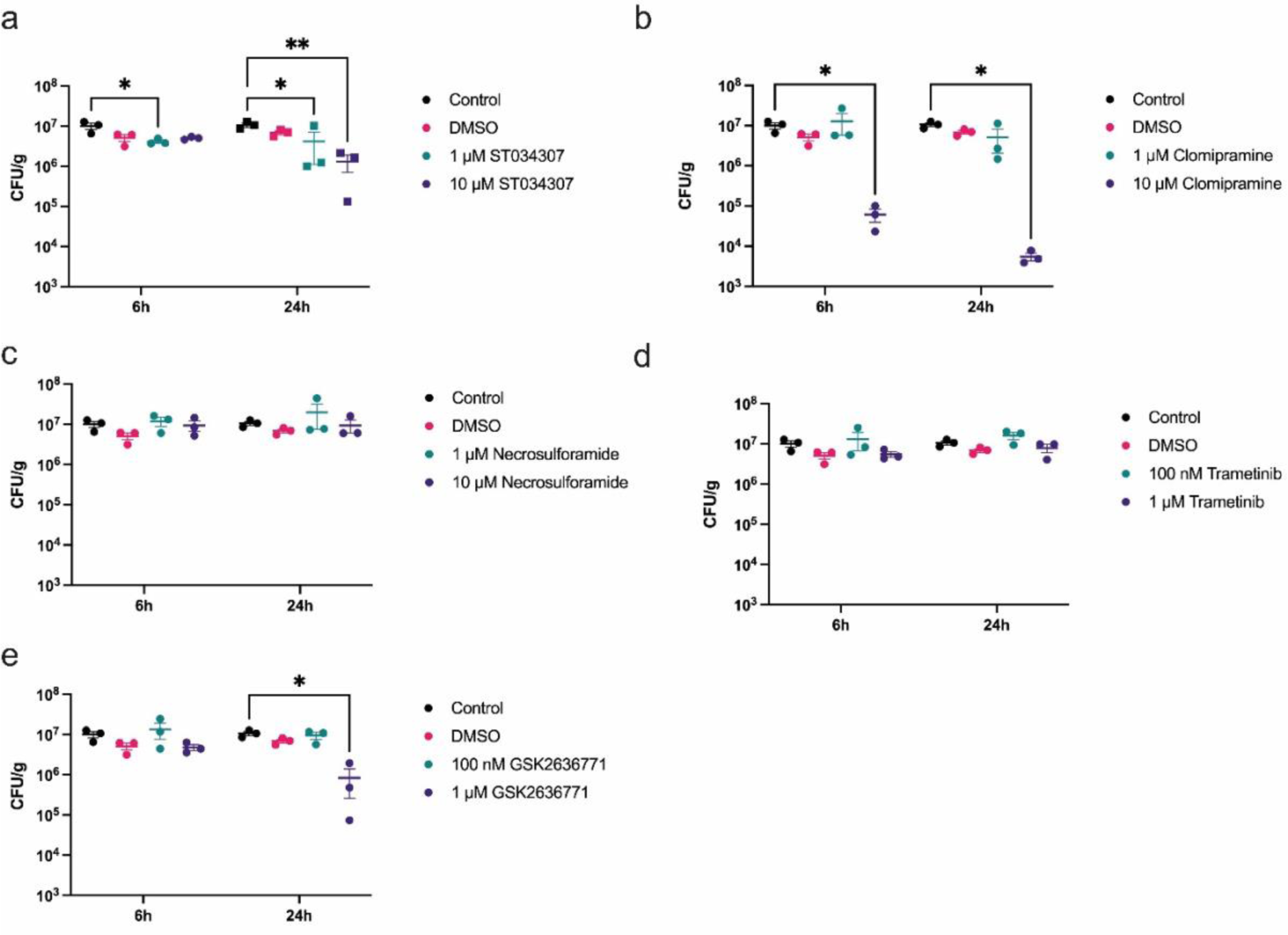
Evaluation of the effects of different chemical inhibitors on intracellular bacterial load in RAW 264.7 cells. RAW 264.7 cells were infected with LF82 for 1 hour followed by treatment with different chemical inhibitors for a further 6 or 24 hpi; ST034307 (a), Clomipramine (b), Necrosulforamide (c), Trametinib (d), GSK2636771 (e). Bacterial recovery is displayed as CFU/g of protein. Data points represent the mean of three technical repeats plus the standard deviation at a timepoint of 6 or 24 hpi. Each treatment was compared to the untreated control group. Statistical significance was determined by two-way ANOVA. *, *P* <0 0.05. **, *P* < 0.01. ***, *P* < 0.0001.

**Figure 8:**
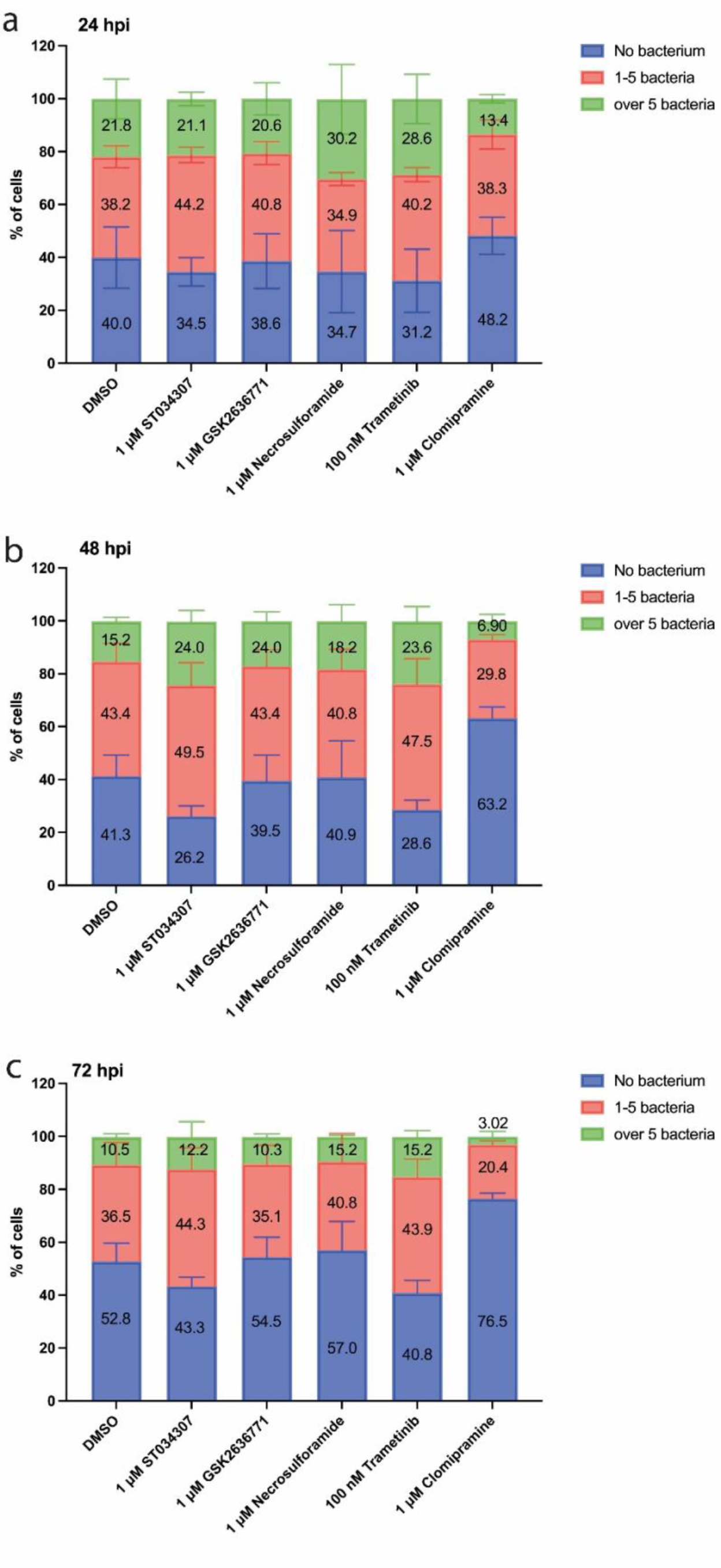
Quantification of intracellular LF82 burden post-inhibitor treatment using imaging flow cytometry. RAW 264.7 cells infected with LF82::*rpsM*GFP were treated with 1 μM ST034307, 1 uM GSK2636771, 1 uM Necrosulforamide, 100 nM Trametinib or 1 uM Clomipramine for 24 hpi (a), 48 hpi (b) and 72 hpi (c). Infected cells treated with DMSO were used as a control. Intracellular LF82::*rpsM*GFP was counted via IFC. The spot count profile separated cells into those with no bacteria, cells containing 1-5 bacteria, or cells containing over 5 bacteria. The sub-populations of a graph represent the mean of three biological repeats. Error bars represent SEM. The number of portions of sub-populations represents the mean of three biological repeats.

**Table 3:**
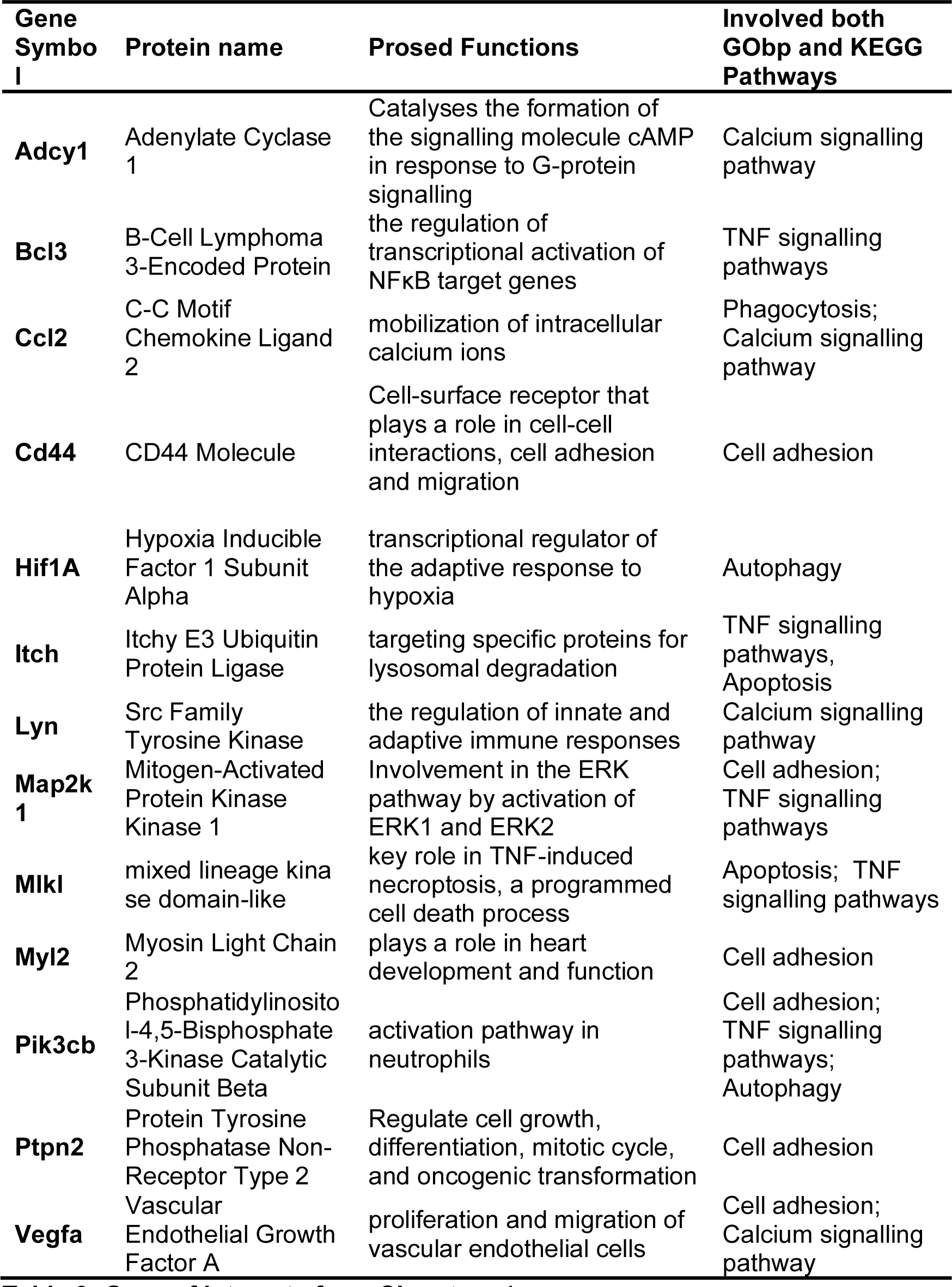
Gene of interests from *Signature 1*.

**Table 4:**
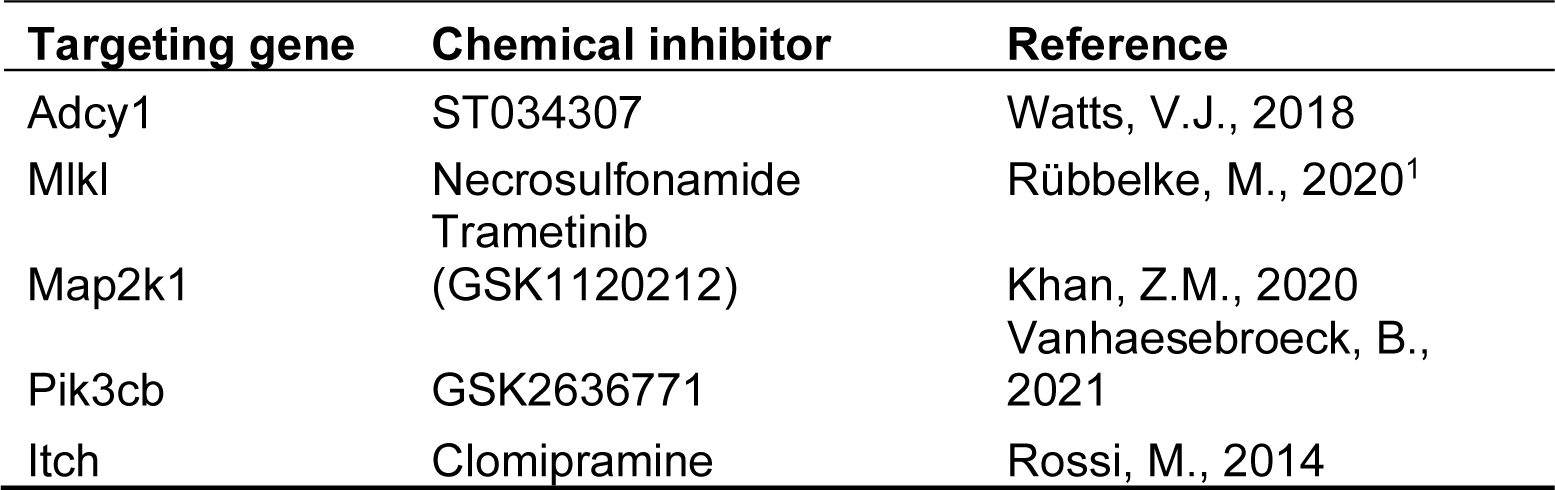
Selected *Signature 1* genes and relevant chemical inhibitors.

Given pathway analysis using *Signature 1* had highlighted a significant role for inflammation and migration of immune cells in response to increasing bacterial burden we next determined any effects of the identified inhibitors on TNFα release by the infected cells.

TNFα levels were determined post-infection and treatment with the inhibitors (Fig. 9). Trametinib significantly inhibited TNFα release by both infected and uninfected cells at both 100 nM and 1 μM with the reduction apparent at early (6 hpi) and later points of infection (24 hpi). While ST034307 inhibited TNFα, the reduction was only apparent at a higher cytotoxic concentrations of the inhibitor (10 μM) in infected cells (Fig. 9b and 9d). Most interestingly however it is noticeable that trametinib, while it significantly reduced TNFα release by infected cells, did not reduce intracellular LF82 burden. Contrastingly clomipramine, which reduced intracellular LF82 burden, had no impact on TNFα release. This disconnect between AIEC infection and cytokine release by immune cells has not previously been described and may offer future opportunities for intervention to prevent inflammation despite bacterial burdens.

**Figure 9:**
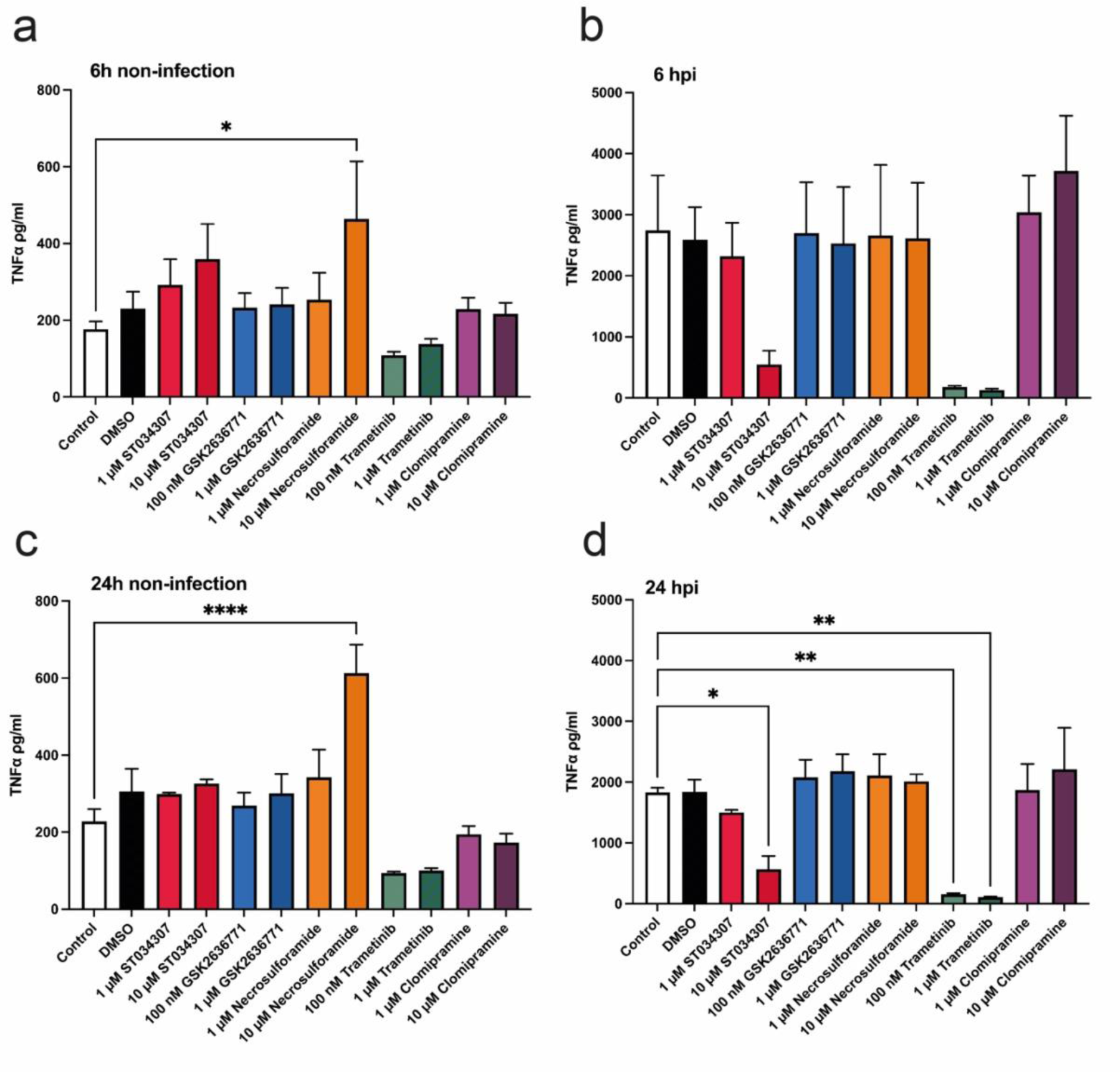
TNFα secretion by RAW 264.7 cells measurement post-inhibitor treatment. RAW 264.7 cells were stimulated overnight by 100 ng/ml LPS. Activated RAW 264,7 cells were then infected with LF82 at MOI of 100 or treated with bacteria-free medium (as uninfected RAW 264.7 cells) for 1 hour. One hour post-infection, infected or uninfected RAW 264.7 cells were washed and treated with different chemical inhibitors at two different concentrations for further indicated times. Infected or uninfected cells in absence of chemical treatment was regarded as a control. Graph (**a**) represents uninfected RAW 264.7 cells that were treated with or without chemical treatments for 6 hpi. (**b**) As described for (**a**) but for infected RAW 264.7 cells. (**c**) Uninfected RAW 264.7 cells were treated with chemical inhibitors for 24 hpi. (**d**) As for (**c**) but for infected RAW 264.7 cells. Statistical significance was determined by one-way ANOVA. *, P <0 0.05. **, P < 0.01. ***, P < 0.0001.

## Discussion

Bacterial infection, both *in vitro* and *in vivo*, results in a heterogenous population of cells, comprising those infected to differing levels by the pathogen, and those that remain uninfected, termed bystander cells. The diversity of outcomes at the cellular level presents a conundrum as regards studying infection, as the mixed population can have an array of bacterial burdens resulting in diverse host and microbial gene expression. This heterogeneity makes interpretation of the host response particularly difficult as it can mask crucial host mediators of infection.

Here we demonstrate this heterogeneity within an LF82-treated well of RAW 264.7 cells *in vitro*. Close to 60% of cells carry intracellular LF82 at 24 hpi, but this results in any subsequent analysis of host gene expression in response to infection including the remaining 40% of cells that are uninfected. Even within the LF82-bearing cells our data demonstrates that two thirds of these cells contain less than 5 bacteria, within only 20% of the total population of cells bearing more than 5 bacteria. Given that intracellular replication in immune cells has been described as a critical phenotypic marker of this pathobiont, the fact that only one fifth of the infected population of cells meet this criterium makes it challenging to study (Bringer *et al*., 2012). Any host transcriptional changes in response to intracellular replication of AIEC will be difficult to pick up in downstream analysis due to being masked by the transcriptional changes in the remaining 80% of cells.

To overcome the challenges of a heterogeneously infected population, here we took an approach of cell sorting based on intracellular bacterial load followed by RNA sequencing. This enabled us to stratify the heterogenous population into distinct population subsets, each with its own characteristics of being uninfected or infected and, if infected, stratified into further sub-populations based on their intracellular LF82 burden. Clearly there were significant differences between cells exposed to LF82 and unexposed and uninfected cells. Surprisingly bystander cells from wells where LF82 was present, but with no intracellular LF82, displayed a phenotypic shift that mirrored that of infected cells, and which was distinct from cells from uninfected wells. Over 400 genes were significantly differentially regulated between both these uninfected populations, with 77 of the DEGs from these bystander cells unique to them and not identified in infected cells from the same well. This indicated that while also responding to LF82 in a manner similar to infected cells, these bystander cells were a unique population in themselves.

Interestingly our analyses also indicated that uninfected bystander cells were directly contributing to inflammation despite not being actively infected with AIEC. It is likely that immune activation of these bystander cells is driven by either contact with bacteria, bacteria-derived molecules and vesicles being shed into the media, or immune cell derived TNFα (Bringer *et al*., 2012; Jung *et al*., 2017; Qu, Zhu and Zhang, 2022). However, the relative contribution of each to bystander cell activation cannot be ascertained from the data generated here, but understanding this could be informative in the context of CD given the importance of TNFα in driving inflammation in CD. If bystander cell activation, and their subsequent contribution to inflammation, was dependent on TNFα, anti-TNFα therapy as used in CD would block activation of these cells. Also as immune cells with a high intracellular bacterial burden are likely to be the primary source of TNFα sparking a subsequent inflammatory cascade, targeting this small population of highly infected cells to remove AIEC would offer most therapeutic benefit.

Differences in cells either exposed to, or infected with, LF82 were further underlined through direct comparison of gene expression amongst these groups. Again, unique DEGs were found for each group with some DEGs common to more than one sub-population within an infected well. A major advantage of our approach was the ability to directly match host gene expression to bacterial load across the stratified sub-populations. Given the depth of data available, and the significant number of DEGs identified, an approach of enrichment analysis whereby signatures of gene expression were correlated to bacterial load was undertaken.

This approach identified DEGs and pathways directly responding to increasing or decreasing intracellular bacterial load. While gene expression may fluctuate due to bacterial load, using signatures of infection across populations allowed us to concentrate on DEGs whose expression was directly related to infectious burden. This approach of identifying signatures of gene expression in response to intracellular infectious load revealed several pathways related to increasing or decreasing bacterial load. Given their likely importance to success of infection we targeted these pathways using chemical inhibitors, selecting target proteins from the significant DEGs within these pathways of interest. This enabled testing their role in mediating both intracellular replication of AIEC and its induction of inflammation.

The targets chosen; Adcy1, Pik3cb, Mlkl, Map2k1 and Itch, each represented a unique pathway in which they displayed the *Signature 1* phenotype of increasing in direct response to bacterial burden within the cell. None of these genes had to date been associated with AIEC infection or used as a target to inhibit bacterial infection, although PIK3cb and MLKL had previously been suggested as targets for therapeutic intervention in IBD, while MAP2K1 has a currently approved kinase inhibitor targeted towards it for IBD treatment (Pierdomenico *et al*., 2014; Bruckner *et al*., 2020; Winkelmann *et al*., 2021). ITCH has been directly implicated in pathogenesis of nucleotide-binding oligomerization domain-containing protein 2 (NOD2) mediated inflammatory disease and it is directly involved in ubiquitination and tagging of host proteins for proteasomal degradation, a system we have previously shown to be exploited during AIEC infection (Dunne *et al*., 2013). Concentrations of each inhibitor used were those previously published in the literature although it was noted that some caused increased cytotoxicity during testing on RAW 264.7 cells, and this was exacerbated by infection in the case of the Adcy1 inhibitor ST034307 (Břehová *et al*., 2021). Inhibition of Adcy1, Mlkl or Pik3cb function had no significant effect on LF82 infection over the time tested, with no reduction in either intracellular bacterial burden or release of inflammatory cytokines. However, the inhibitor of Itch, trametinib, alongside the inhibitor of Map2K1, clomipramine, generated intriguing results. While clomipramine significantly reduced both intracellular burden of LF82 and the number of cells infected with LF82, trametinib significantly inhibited TNFα release. Intriguingly in the case of both inhibitors, they decoupled intracellular proliferation and cytokine release which have been shown to be interdependent during AIEC infection (Bringer *et al*., 2012; Douadi *et al*., 2022). Kinase inhibitors such as trametinib can block cell proliferation, arrest the cell cycle and induce cell death as well as blocking extracellular signal-regulated kinase (ERK) signalling, which plays a role in cytokine secretion during AIEC infection (Hedl and Abraham, 2012; Hoffner, MSN, ANP-BC, AOCNP and Benchich, MSN, NP-C, AOCNP, 2018). Given proliferation of infected cells is unlikely as cell cycle arrest is already occurring during LF82 infection based on the *Signature 2* pathways identified, the reduction in TNFα secretion observed is likely due to trametinib interruption of signalling pathways, such as that controlled by ERK, upstream of TNFα release.

The mechanism of action of clomipramine in the context of LF82 infection was more challenging to interpret. Used to treat obsessive compulsive disorder, clomipramine effects are likely mediated through reducing re-uptake of norepinephrine and serotonin. However, it has recently been used to treat both viral and parasitic infections with a suggested mechanism of action related to its effects on lysosomal pH undermining viral protease efficacy (Vater *et al*., 2017; Nobile *et al*., 2020; Strauss *et al*., 2021; Khan *et al*., 2022). With lysosomal defence integral to combatting AIEC infection this may explain the phenotype observed here (Spalinger *et al*., 2022). Clomipramine effect on intracellular LF82 replication was clear cut, significantly reducing both the intracellular bacterial load within cells and the number of cells carrying bacteria. Most strikingly, given previous work describing how LF82 intramacrophage replication and TNFα release were intertwined, this reduction in LF82 numbers showed no effect on TNFα release. This disconnect between TNFα mediated inflammation and AIEC intracellular replication, which to now have described as mutually dependent, may help in unravelling the complex host-AIEC relationship.

The data presented here therefore clearly demonstrates that stratifying infected populations of immune cells into distinct sub-populations based on their bacterial load can reveal new therapeutic targets in infection. Here this approach has shed light on tackling a crucial population of inflammatory immune cells in CD, those heavily infected with AIEC. This targeted approach is relatively simplistic but clearly showed promise, with chemical inhibition of target genes either blocking intracellular replication or reducing secretion of TNFα. This is the first time an approach has specifically targeted and been effective against heavily AIEC infected immune cells.

## Acknowledgements

The authors gratefully acknowledge the Sii-Flow Cytometry Core Facility at the University of Glasgow for their support & assistance in this work.

---

**Figure S1:**
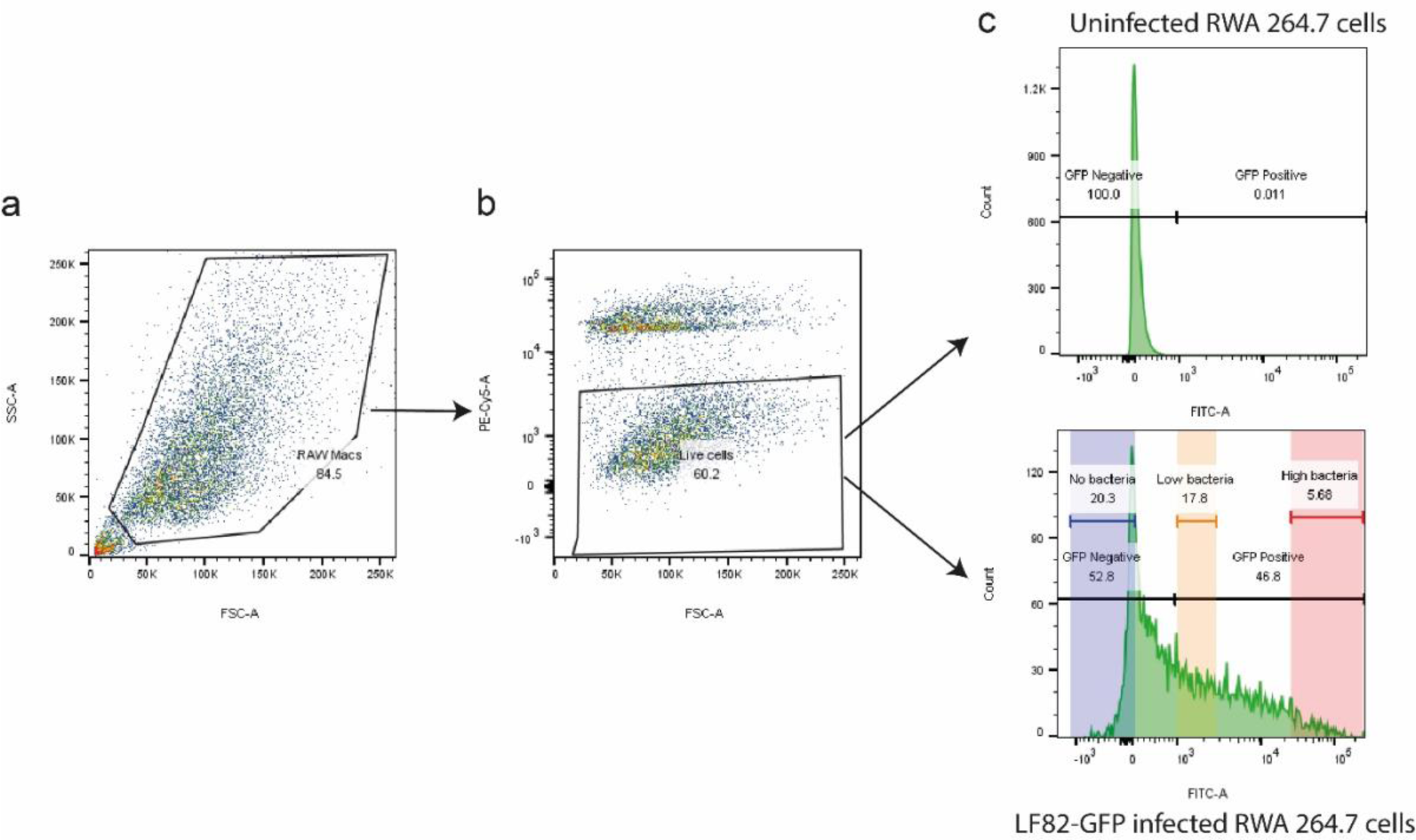
Gating strategy for isolation of LF82::*rpsM*GFP infected RAW 264.7 cells for three different populations (*No*, *Low* and *High*). (**a**) For the isolation of a highly pure RAW 264.7 population, cells were gated on their forward scatter area (FSC-A) and side scatter area (SSC-A), excluding debris from the live gate. (**b**) Dead cells were further excluded based on FSC-A versus the intensity of 7AAD. (**c**) This was followed by gating out three sub-populations of living RAW 264.7 cells according to GFP intensity, resulting in sorting final three populations including cells with no bacterial burden, low bacterial burden and high bacterial burden.

**Figure S2:**
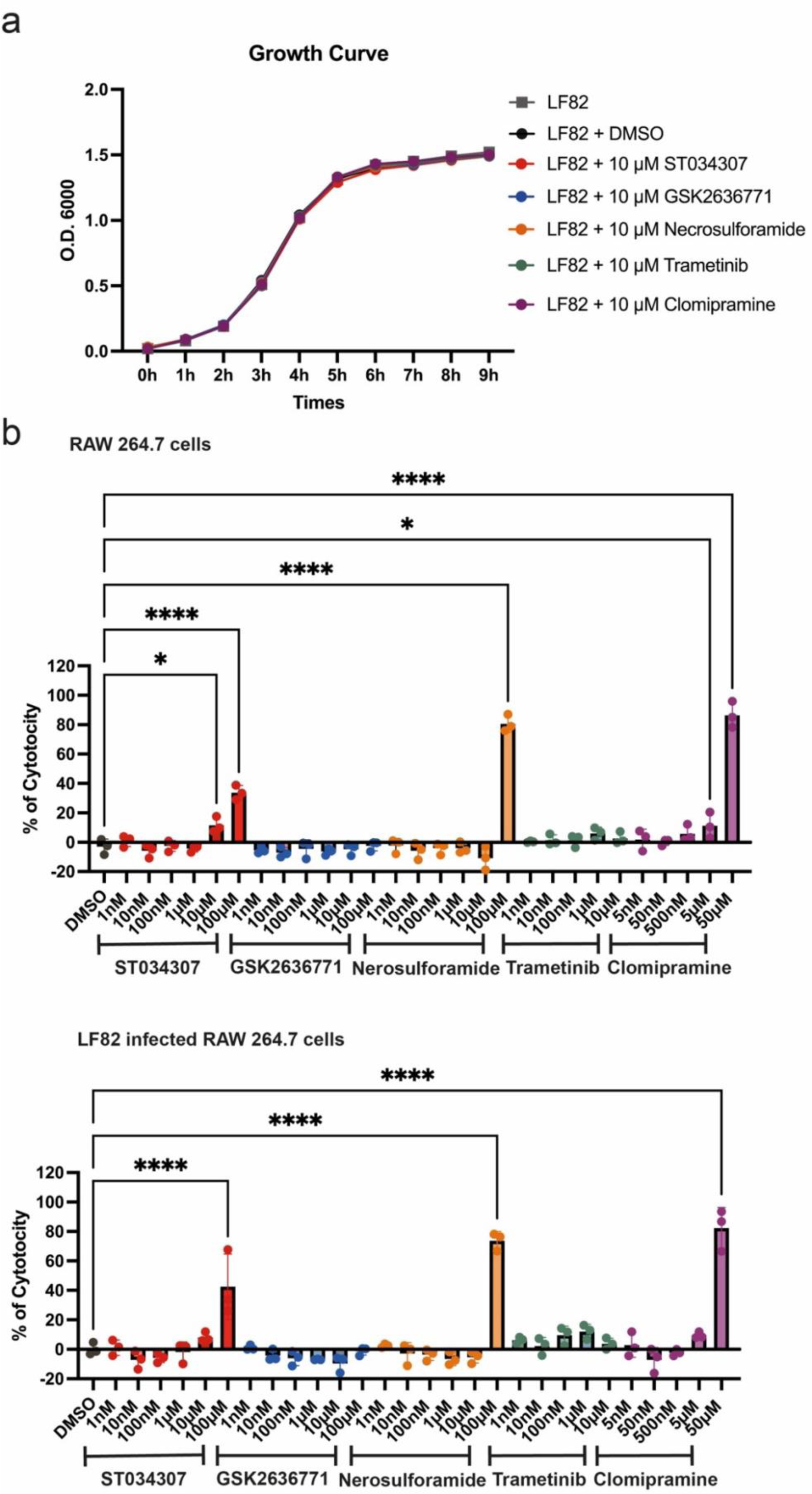
Effects of the different chemical inhibitors on LF82 growth and cytotoxicity to RAW 264.7 cells. (**a**) Growth curve of LF82 after treatment with DMSO or the five chemical inhibitors (10 μM ST034307, 10 μM GSK2636771, 10 μM Necrosulforamide, 10 μM Trametinib or 10 μM Clomipramine). (**b**) Lactate dehydrogenase (LDH) cytotoxicity assay was undertaken for uninfected or LF82 infected RAW 264.7 cells treated with different concentrations of chemical inhibitors. DMSO, a diluent for the inhibitors, was used as a control. Experimental groups were compared to the control. Statistical significance was determined by one-way ANOVA. Statistical significance was determined by one-way ANOVA. *, *P* <0 0.05. **, *P* < 0.01. ***, *P* < 0.0001.

## Bibliography

1. Anders, S., Pyl, P. T. and Huber, W. (2015) ‘HTSeq--a Python framework to work with high-throughput sequencing data’, *Bioinformatics (Oxford*, England). Bioinformatics, 31(2), pp. 166–169. doi: 10.1093/BIOINFORMATICS/BTU638.

2. Bassi, A. et al. (2004) ‘Cost of illness of inflammatory bowel disease in the UK: a single centre retrospective study’, Gut. BMJ Publishing Group, 53(10), pp. 1471–1478. doi: 10.1136/GUT.2004.041616.

3. Bolger, A. M., Lohse, M. and Usadel, B. (2014) ‘Trimmomatic: a flexible trimmer for Illumina sequence data’, *Bioinformatics (Oxford*, England). Bioinformatics, 30(15), pp. 2114–2120. doi: 10.1093/BIOINFORMATICS/BTU170.

4. Boucher, D. and Barnich, N. (2022) ‘Phage Therapy Against Adherent-invasive E. coli: Towards a Promising Treatment of Crohn’s Disease Patients?’, Journal of Crohn’s & colitis. NLM (Medline), 16(10), pp. 1509–1510. doi: 10.1093/ecco-jcc/jjac070.

5. Břehová, P. et al. (2021) ‘Acyclic nucleoside phosphonates with 2-aminothiazole base as inhibitors of bacterial and mammalian adenylate cyclases’, European Journal of Medicinal Chemistry, 222. doi: 10.1016/j.ejmech.2021.113581.

6. Bringer, M. A. et al. (2012) ‘Replication of Crohn’s disease-associated AIEC within macrophages is dependent on TNF-α secretion’, *Laboratory Investigation*. Lab Invest, 92(3), pp. 411–419. doi: 10.1038/labinvest.2011.156.

7. Bruckner, R. S. et al. (2020) ‘DOP86 An IFN-STAT1-MLKL axis drives programmed necrosis of Paneth cells in Crohn’s ileitis’, Journal of Crohn’s and Colitis. Oxford Academic, 14(Supplement_1), pp. S125–S126. doi: 10.1093/ECCO-JCC/JJZ203.125.

8. Cho, Y. H. et al. (2022) ‘Inflammatory bowel disease-associated adherent-invasive Escherichia coli have elevated host-defense peptide resistance’, *FEMS microbiology letters*. FEMS Microbiol Lett, 369(1). doi: 10.1093/FEMSLE/FNAC098.

9. Cole, J. J. et al. (2021) ‘Searchlight: automated bulk RNA-seq exploration and visualisation using dynamically generated R scripts’, BMC Bioinformatics. BioMed Central Ltd, 22(1), pp. 1–21. doi: 10.1186/S12859-021-04321-2/TABLES/1.

10. Darfeuille-Michaud, A. et al. (1998) ‘Presence of adherent *Escherichia coli* strains in ileal mucosa of patients with Crohn’s disease.’, Gastroenterology, 115(6), pp. 1405–1413.

11. Douadi, C. et al. (2022) ‘Anti-TNF Agents Restrict Adherent-invasive Escherichia coli Replication Within Macrophages Through Modulation of Chitinase 3-like 1 in Patients with Crohn’s Disease’, *Journal of Crohn’s & colitis*. J Crohns Colitis, 16(7), pp. 1140–1150. doi: 10.1093/ECCO-JCC/JJAB236.

12. Dunne, K. A. et al. (2013) ‘Increased S-nitrosylation and proteasomal degradation of caspase-3 during infection contribute to the persistence of adherent invasive Escherichia coli (AIEC) in immune cells.’, PloS one, 8(7), p. e68386. doi: 10.1371/journal.pone.0068386.

13. Fleige, S. and Pfaffl, M. W. (2006) ‘RNA integrity and the effect on the real-time qRT-PCR performance’, *Molecular aspects of medicine*. Mol Aspects Med, 27(2–3), pp. 126–139. doi: 10.1016/J.MAM.2005.12.003.

14. Gerner, R. R. et al. (2022) ‘Siderophore Immunization Restricted Colonization of Adherent- Invasive Escherichia coli and Ameliorated Experimental Colitis’, *mBio*. mBio, 13(5). doi: 10.1128/MBIO.02184-22.

15. Hedl, M. and Abraham, C. (2012) ‘Nod2-Induced Autocrine Interleukin-1 Alters Signaling by ERK and p38 to Differentially Regulate Secretion of Inflammatory Cytokines’, Gastroenterology. NIH Public Access, 143(6), p. 1530. doi: 10.1053/J.GASTRO.2012.08.048.

16. Hoffner, MSN, ANP-BC, AOCNP, B. and Benchich, MSN, NP-C, AOCNP, K. (2018) ‘Trametinib: A Targeted Therapy in Metastatic Melanoma’, Journal of the Advanced Practitioner in Oncology. Harborside Press, 9(7), p. 741. doi: 10.6004/jadpro.2018.9.7.5.

17. Jung, A. L. et al. (2017) ‘Legionella pneumophila infection activates bystander cells differentially by bacterial and host cell vesicles’, Scientific Reports. Springer US, 7(1), pp. 1–11. doi: 10.1038/s41598-017-06443-1.

18. Khan, E. et al. (2022) ‘Anti-depressants and COVID-19: A New Ray of Hope’, *Psychiatria Danubina*. Psychiatr Danub, 34(1), pp. 171–173. doi: 10.24869/PSYD.2022.171.

19. Kim, D. et al. (2019) ‘Graph-based genome alignment and genotyping with HISAT2 and HISAT-genotype’, *Nature biotechnology*. Nat Biotechnol, 37(8), pp. 907–915. doi: 10.1038/S41587-019-0201-4.

20. Li, X. et al. (2022) ‘Inhibition of proline tyrosine kinase 2 (Pyk2) phosphorylation during adherent-invasive Escherichia coli infection inhibits intra-macrophage replication and inflammatory cytokine release’, *bioRxiv*. Cold Spring Harbor Laboratory, p.2022.11.14.516411. doi: 10.1101/2022.11.14.516411.

21. Martinez-Medina, M. et al. (2009) ‘Molecular diversity of *Escherichia coli* in the human gut: New ecological evidence supporting the role of adherent-invasive *E. coli* (AIEC) in Crohn’s disease’, Inflammatory Bowel Diseases, 15(6), pp. 872–882. doi: 10.1002/ibd.20860.

22. Meconi, S. et al. (2007) ‘Adherent-invasive Escherichia coli isolated from Crohn’s disease patients induce granulomas in vitro’, Cellular Microbiology, 9(5), pp. 1252–1261. doi: 10.1111/j.1462-5822.2006.00868.x.

23. Nadalian, B. et al. (2021) ‘Prevalence of the pathobiont adherent-invasive Escherichia coli and inflammatory bowel disease: a systematic review and meta-analysis’, *Journal of gastroenterology and hepatology*. J Gastroenterol Hepatol, 36(4), pp. 852–863. doi: 10.1111/JGH.15260.

24. Nobile, B. et al. (2020) ‘Clomipramine Could Be Useful in Preventing Neurological Complications of SARS-CoV-2 Infection’, *Journal of neuroimmune pharmacology : the official journal of the Society on NeuroImmune Pharmacology*. J Neuroimmune Pharmacol, 15(3), pp. 347–348. doi: 10.1007/S11481-020-09939-2.

25. Ormsby, M. J. et al. (2019) ‘Inflammation associated ethanolamine facilitates infection by Crohn’s disease-linked adherent-invasive Escherichia coli’, EBioMedicine. Elsevier B.V., 43, pp. 325–332. doi: 10.1016/j.ebiom.2019.03.071.

26. Ormsby, M. J. et al. (2020) ‘Propionic Acid Promotes the Virulent Phenotype of Crohn’s Disease-Associated Adherent-Invasive Escherichia coli’, Cell Reports, 30(7). doi: 10.1016/j.celrep.2020.01.078.

27. Pierdomenico, M. et al. (2014) ‘Necroptosis is active in children with inflammatory bowel disease and contributes to heighten intestinal inflammation’, *The American journal of gastroenterology*. Am J Gastroenterol, 109(2), pp. 279–287. doi: 10.1038/AJG.2013.403.

28. Qu, M., Zhu, H. and Zhang, X. (2022) ‘Extracellular vesicle-mediated regulation of macrophage polarization in bacterial infections’, Frontiers in Microbiology, 13. doi: 10.3389/fmicb.2022.1039040.

29. Rao, B. B. et al. (2017) ‘The Cost of Crohn’s disease: varied healthcare expenditure patterns across distinct disease trajectories’, Inflammatory bowel diseases. NIH Public Access, 23(1), p. 107. doi: 10.1097/MIB.0000000000000977.

30. Spalinger, M. R. et al. (2022) ‘Original research: Autoimmune susceptibility gene PTPN2 is required for clearance of adherent-invasive Escherichia coli by integrating bacterial uptake and lysosomal defence’, Gut. BMJ Publishing Group, 71(1), p. 89. doi: 10.1136/GUTJNL-2020-323636.

31. Strauss, M. et al. (2021) ‘Differential tissue distribution of discrete typing units after drug combination therapy in experimental Trypanosoma cruzi mixed infection’, *Parasitology*. Parasitology, 148(13), pp. 1595–1601. doi: 10.1017/S0031182021001281.

32. Sugihara, K. et al. (2022) ‘Mucolytic bacteria license pathobionts to acquire host-derived nutrients during dietary nutrient restriction’, *Cell reports*. Cell Rep, 40(3). doi: 10.1016/J.CELREP.2022.111093.

33. Titécat, M. et al. (2022) ‘Safety and Efficacy of an AIEC-targeted Bacteriophage Cocktail in a Mice Colitis Model’, *Journal of Crohn’s & colitis*. J Crohns Colitis, 16(10), pp. 1617–1627. doi: 10.1093/ECCO-JCC/JJAC064.

34. Vater, M. et al. (2017) ‘New insights into the intracellular distribution pattern of cationic amphiphilic drugs’, Scientific Reports. Nature Publishing Group, 7. doi: 10.1038/SREP44277.

35. Winkelmann, P. et al. (2021) ‘The PI3K pathway as a therapeutic intervention point in inflammatory bowel disease’, *Immunity, inflammation and disease*. Immun Inflamm Dis, 9(3), pp. 804–818. doi: 10.1002/IID3.435.

